# Evidence for the role of selection for reproductively advantageous alleles in human aging

**DOI:** 10.1101/2023.04.13.536806

**Authors:** Erping Long, Jianzhi Zhang

## Abstract

The antagonistic pleiotropy hypothesis posits that natural selection for pleiotropic mutations that confer earlier or more reproduction but impair the post-reproductive life causes aging. This hypothesis of the evolutionary origin of aging is supported by case studies but lacks unambiguous genomic evidence. Here we genomically test this hypothesis using the genotypes, reproductive phenotypes, and death registry of 276,406 UK Biobank participants. We observe a strong, negative genetic correlation between reproductive traits and lifespan. Individuals with higher polygenetic scores for reproduction (*PGS*_R_) have lower survivorships to age 76 (*SV*_76_), and *PGS*_R_ increased over birth cohorts from 1940 to 1969. Similar trends are found from individual genetic variants examined. *PGS*_R_ and *SV*_76_ remain negatively correlated upon the control of the offspring number, revealing horizontal pleiotropy between reproduction and lifespan. Intriguingly, regardless of *PGS*_R_, having two children maximizes *SV*_76_. These and other findings strongly support the antagonistic pleiotropy hypothesis of aging in humans.

## INTRODUCTION

Aging or senescence refers to a gradual deterioration of bodily functions that manifests as an increase in the death rate with age after sexual maturity. The prevalence of aging across multicellular organisms has been an evolutionary puzzle because natural selection should favor mutations conferring an extended reproductive lifespan^1^. In 1957, George Williams proposed that mutations contributing to aging could be positively selected if they are advantageous early in life such as promoting development or reproduction that leads to earlier reproduction or more offspring^2^. This antagonistic pleiotropy hypothesis has become one of the leading theories of the evolutionary origin of aging (see Discussion for other theories). Nonetheless, despite observations in life-history evolution of tradeoffs consistent with the antagonistic pleiotropy hypothesis^3,4^, the involvement of mutational pleiotropy is usually difficult to establish^5^. Experimental studies of model organisms have identified cases of antagonistic pleiotropy between reproduction and lifespan^5–8^, but how frequently pleiotropy occurs between reproduction and longevity and how common such pleiotropy is antagonistic as opposed to concordant are unknown at the genomic scale. Further complicating the situation is the fact that phenotypes measured in the lab can often deviate from those measured in nature^5^. In humans, cases consistent with the antagonistic pleiotropy hypothesis are known^9,10^. For example, genetic variants associated with coronary artery disease tend to be associated with reproductive success^9^. However, evidence for the antagonistic pleiotropy hypothesis in humans have been contested^11–14^, and the hypothesis still lacks a genomic test. Furthermore, it is unclear whether mutations contributing to aging were selectively favored and whether the selection arose from their benefits earlier in life.

One of the primary reasons why testing the antagonistic pleiotropy hypothesis is difficult is that life-history traits tend to be influenced by many small-effect genetic variants^15,16^. The UK Biobank has collected the genotypes and various phenotypes of about 0.5 million participants^17^, offering an unprecedented opportunity for testing the antagonistic pleiotropy hypothesis in humans. In particular, the data include multi-dimensional measures of human reproduction from sexual maturation (e.g., menarche) to reproduction (e.g., number of offspring) and infertility (e.g., sexual dysfunction) and include exact death dates from the national death registry^18^. Such phenotypic data along with genotypic data allow assessing the relationship between reproduction and lifespan at the genomic scale as well as at the level of individual segregating variants.

Furthermore, functional genomic data (e.g., expression quantitative trait locus, eQTL) can be mined to uncover the putative pathways and mechanisms underlying any reproduction-lifespan relationship^19,20^. Taking advantage of these resources, we performed multiple analyses outlined in **Fig. 1** to address the following questions about the antagonistic pleiotropy hypothesis. First, do genetic variants influencing reproduction more likely affect lifespan than expected by chance? Second, is the pleiotropy between reproduction and lifespan largely antagonistic? Third, are pleiotropic mutations promoting reproduction but causing aging favored by natural selection? Fourth, what are the potential mechanisms linking reproduction and aging?

**Figure 1.**
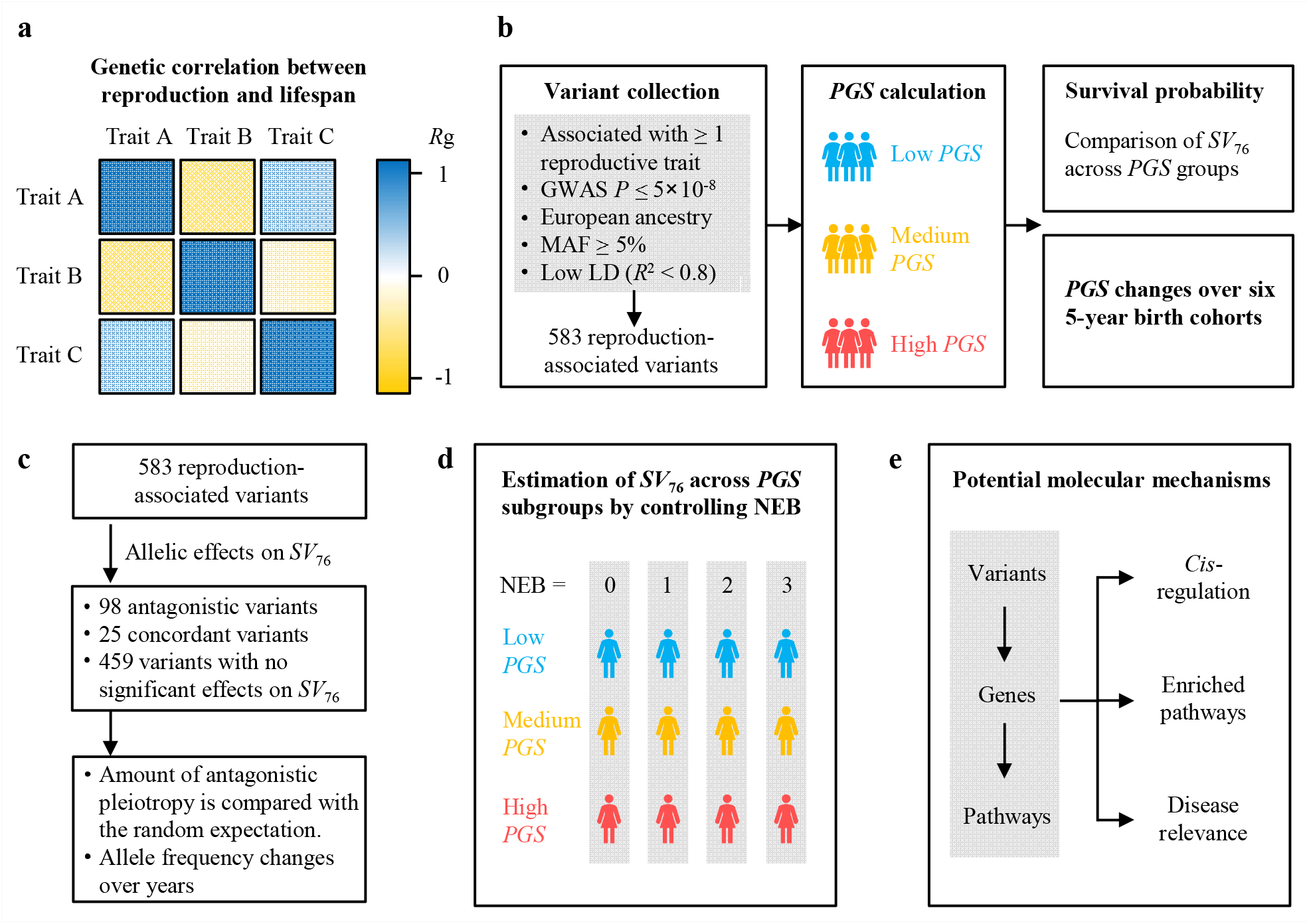
Schematics summarizing the approaches and analyses used in the present study. **a**, Genetic correlation between reproduction and lifespan identified from a list of previously computed genetic correlations for heritable UK Biobank traits. Available traits include three reproductive traits—negative age at first birth (nAFB), negative age at first sex (nAFS), and number of children fathered (NCF), as well as two parental lifespan traits— father’s age at death and mother’s age at death. **b**, A total of 583 reproduction-associated variants are collected, and polygenic scores (PGSs) are calculated for four reproductive traits: (1) nAFB, (2) nAFS, (3) negative age at menarche (nAMC), and (4) age at menopause (AMP). The probability of survival to age 76 (*SV*_76_) is compared across equal-size groups of individuals with relatively low, medium, and high PGS for each of the four reproductive traits. The potential change of PGS over six 5-year birth cohorts from 1940 to 1969 is investigated. **c**, Effects of individual variants on lifespan are estimated from *SV*_76_. The observed amount of antagonistic pleiotropy is compared with the random expectation. Potential allele frequency changes over six 5-year birth cohorts from 1940 to 1969 are examined. **d**, Potential horizontal pleiotropy between reproduction and lifespan is explored by controlling the number of children ever born (NEB = 0, 1, 2, or 3). **e**, Potential molecular mechanisms of antagonistic pleiotropy are explored by surveying the *cis*-regulatory activity of antagonistically pleiotropic variants, their target genes, enriched pathways, and relevance to human diseases.

## RESULTS

### Negative genetic correlation between reproduction and lifespan

The genetic correlation between two phenotypic traits is the proportion of variance that the two traits share due to genetic causes and is a measure of the contribution of pleiotropy to the covariation of the traits^21^. Because the antagonistic pleiotropy hypothesis of aging posits that the same mutations antagonistically impact reproduction and lifespan, the hypothesis predicts a negative genetic correlation between reproduction and lifespan. Because the lifespans of most UK Biobank participants are unknown (as they are still living), we examined the genetic correlation between the reproductive traits of UK Biobank participants and the lifespans of their parents by searching the genetic correlations between heritable traits in UK Biobank previously computed by the Neale group^22^ (see Materials and Methods). Note that both the number of offspring and timing of reproduction impact a person’s fitness, because earlier reproduction means a shorter generation time and thereby a higher fitness even when the number of offspring is given. We therefore focused on three reproductive traits available in the list—negative age at first birth (nAFB), negative age at first sex (nAFS), and number of children fathered (NCF); larger values of these traits correspond to higher reproduction (i.e., earlier reproduction and/or more offspring). Note that nAFB is a more accurate measure of the earliest reproduction of a participant than nAFS, but nAFB was measured for female UK Biobank participants only.

Hence, we considered both nAFB and nAFS here, the latter of which was measured for both male and female participants. We examined two parental lifespan traits—father’s age at death and mother’s age at death. Reproduction and parental lifespan show a significant negative genetic correlation in all pairwise comparisons between the three reproductive traits and two lifespan traits (**Table 1**), supporting the antagonistic pleiotropy hypothesis. Because the UK Biobank also recorded the number of full brothers and that of full sisters of each participant, we further examined the genetic correlation between a participant’s parental reproduction and parental lifespan. A significant genetic correlation was observed (**Data S1**), further supporting the antagonistic pleiotropy hypothesis.

**Table 1.** Genetic correlation between reproduction and parental lifespan.

| **Genetic correlation coefficient $R_g$** | **nAFS** | **nAFB** | **NCF** | **Father's age at death** | **Mother's age at death** |
| --- | --- | --- | --- | --- | --- |
| **Negative age at first sex (nAFS)** | 1 | 0.84 | 0.48 | -0.59 | -0.75 |
| **Negative age at first birth (nAFB)** | 0.84 | 1 | 0.45 | -0.66 | -0.80 |
| **No. of children fathered (NCF)** | 0.48 | 0.45 | 1 | -0.29 | -0.33 |
| **Father's age at death** | -0.59 | -0.66 | -0.29 | 1 | 0.91 |
| **Mother's age at death** | -0.75 | -0.80 | -0.33 | 0.91 | 1 |

The above $R_g$ values are from http://www.nealelab.is/blog/2019/10/10/genetic-correlation-results-for-heritable-phenotypes-in-the-uk-biobank and are all significant at $P < 10^{-7}$ except for that between NCF and father's age at death ( $P = 1.4 \times 10^{-5}$ ) and that between NCF and mother's age at death ( $P = 2.0 \times 10^{-5}$ ).

In addition to biological pleiotropy (one mutation affecting multiple traits) and statistical pleiotropy (linked mutations affecting multiple traits), both of which are relevant to the antagonistic pleiotropy hypothesis, genetic correlation could also be caused by two factors unrelated to the antagonistic pleiotropy hypothesis. The first is population stratification and second is cross-trait assortative mating^23^. Our above results should not be affected by population stratification because it had been controlled in genome-wide association analysis (GWAS) based on which genetic correlations were computed^22^. There should be no assortative mating between reproductive traits and lifespan because lifespan is unknown at the time of mating.

### Higher polygenic scores for reproduction predict lower probabilities of survival to age 76

Although the lifespans of many UK Biobank participants are unknown, the deaths of some participants allow reliable estimation of the probability of survival to various ages (up to the age of 76 years), providing an opportunity to assess the potential genetic impact of reproduction on the lifespan. We focused on 276,406 participants of British ancestry with no kinship to other participants in the database (see Materials and Methods) and considered 891 genetic variants reported to be associated with at least one reproductive trait with genome-wide significance (*P* ≤ 5×10^−8^)^24^. After the removal of variants with high linkage disequilibrium (*R*^2^ > 0.8; the variant with a relatively high *P* value is excluded) and those with low minor allele frequencies (< 0.05), 583 genetic variants remained, which are associated with the following reproductive traits: (i) nAFB, (ii) nAFS, (iii) negative age at menarche (nAMC), (iv) age at menopause (AMP), (v) number of children ever born (NEB), which is the number of children fathered (NCF) for a male or number of children mothered (NCM) for a female, and (vi) polycystic ovary syndrome (PCOS) (**Data S2**; see Materials and Methods).

An individual’s polygenic score (PGS) of a trait reflects the individual’s estimated genetic predisposition for the trait; it measures the individual’s likelihood of having the trait based exclusively on genetics without taking environmental factors into account. To assess the aggregated effect of the above reproduction-associated genetic variants, we computed the PGSs of four reproduction traits each associated with at least 50 variants: nAFB, nAFS, nAMC, and AMP (see Materials and Methods). nAMC, nAFS, and nAFB measure the onset of reproduction, while AMP measures the end of reproduction, so together they inform the timing and potential amount of reproduction. NEB and PCOS are not considered here because they each have fewer than 50 variants. The survival rate per year from the age of 40 to 76 years and the cumulative survival probability for homozygotes of each variant were calculated (see Materials and Methods). The log-rank method was used to test the difference in survivorship between two genotypes. For example, individuals ranked in the top third in PGS for nAFB (*PGS*_nAFB_) have a significantly lower probability of survival to age 76 (*SV*_76_ = 0.800) than that of individuals ranked in the bottom third in *PGS*_nAFB_ (*SV*_76_ = 0.839) (*P* = 3.5×10^−4^; **Fig. 2a**), supporting the antagonistic pleiotropy hypothesis. Qualitatively similar results were observed when the PGS for each of the other three reproductive traits was examined (**Fig. 2b, Fig. S1**).

**Figure 2.**
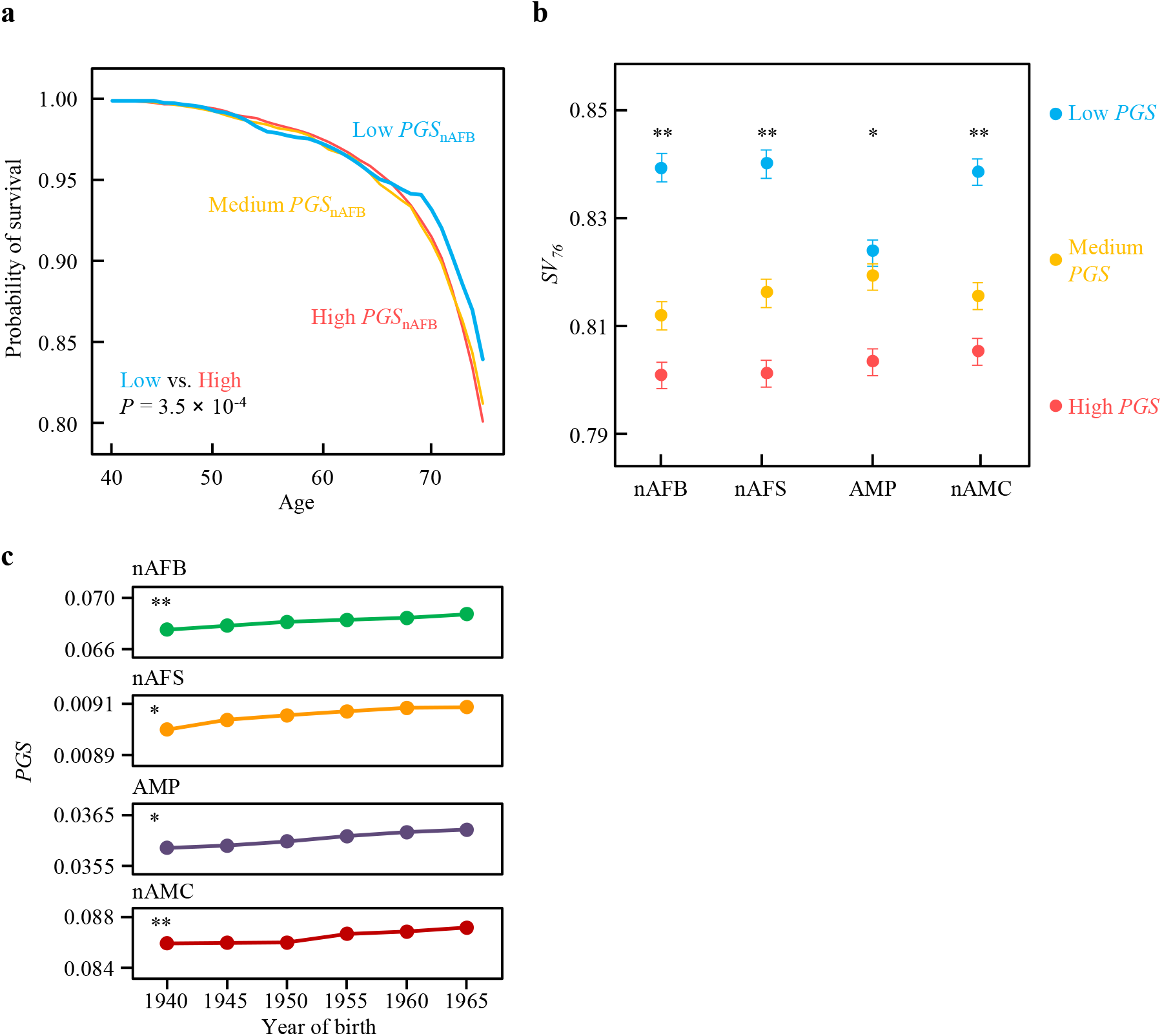
Higher polygenic scores (PGSs) for reproduction predict lower probabilities of survival to the age of 76 years (*SV*_76_). **a**, *SV*_76_ of three equal-size groups of individuals with relatively low, medium, and high PGS for negative age at first birth (*PGS*_nAFB_). *SV*_76_ = 0.839, 0.811, and 0.800 for relatively low, medium, and high *PGS*_nAFB_ groups, respectively. Plots of the other three reproductive traits are in Fig. S1. **b**, *SV*_76_ of three equal-size groups of individuals with relatively low, medium, and high PGS for each of the four reproductive traits (nAFB; nAFS, negative age at first sex; AMP, age at menopause; and nAMC, negative age at menarche). Error bars show 95% confidence intervals calculated by Clopper-Pearson exact methods. *, Log-rank *P* < 0.05; **, *P* < 0.001 between the relatively low and high PGS groups. **c**, Mean PGS for each reproductive trait inferred from genotypes of each of the six 5-year birth cohorts. Error bars showing standard errors are too small to see. *\*, P* < 0.05 and **, *P* < 0.001, linear correlation between mid-point birth year of a cohort and PGS.

Dividing the 276,406 individuals into six 5-year birth cohorts (1940-1944, 1945-1949, …, and 1965-1969), we investigated the change of the PGS of each of the four reproductive traits over birth cohorts. As shown in **Fig. 2c**, for each of these traits, PGS steadily increased over time, presumably a result of natural selection for higher reproduction (see below for the analysis of individual variants that controls potential confounding factors).

Together, the analyses of PGS for reproduction provide evidence for (i) antagonistic pleiotropy between reproduction and lifespan and (ii) natural selection for higher reproduction (presumably to the detriment of lifespan), supporting the antagonistic pleiotropy hypothesis of the origin of aging.

### Genetic variants exhibiting antagonistic pleiotropy between reproduction and lifespan

We further tested the antagonistic pleiotropy hypothesis by identifying genetic variants associated with both reproduction and lifespan. Based on the directions of significant effects of an allele on reproduction and lifespan (see Materials and Methods), we inferred whether a variant shows antagonistic pleiotropy (**Fig. 3a**), concordant pleiotropy (**Fig. 3b**), or no pleiotropy. Among the 583 reproduction-associated variants, 123 variants have significant effects on *SV*_76_ (*FDR* < 0.05), and antagonistic pleiotropy cases significantly outnumber concordant pleiotropy cases (98 antagonistic vs. 25 concordant pleiotropy cases, *P* < 0.00001, two-tailed binomial test; **Fig. 3c, Data S3**). As a negative control, we sampled 100 sets of 583 random variants with matched allele frequencies (± 1%) and recombination rates (± 0.05 cM/Mb), but none of these sets contained ≥ 123 variants with significant effects on *SV*_76_. There is also no significant bias toward antagonistic pleiotropy for these control variants (on average 13 antagonistically vs. 12 concordantly pleiotropic variants per set if the matched allele is considered beneficial to reproduction). Therefore, compared with random polymorphisms, those impacting reproduction are 4.9 times as likely to impact lifespan and 7.5 times as likely to impact lifespan antagonistically.

**Figure 3.**
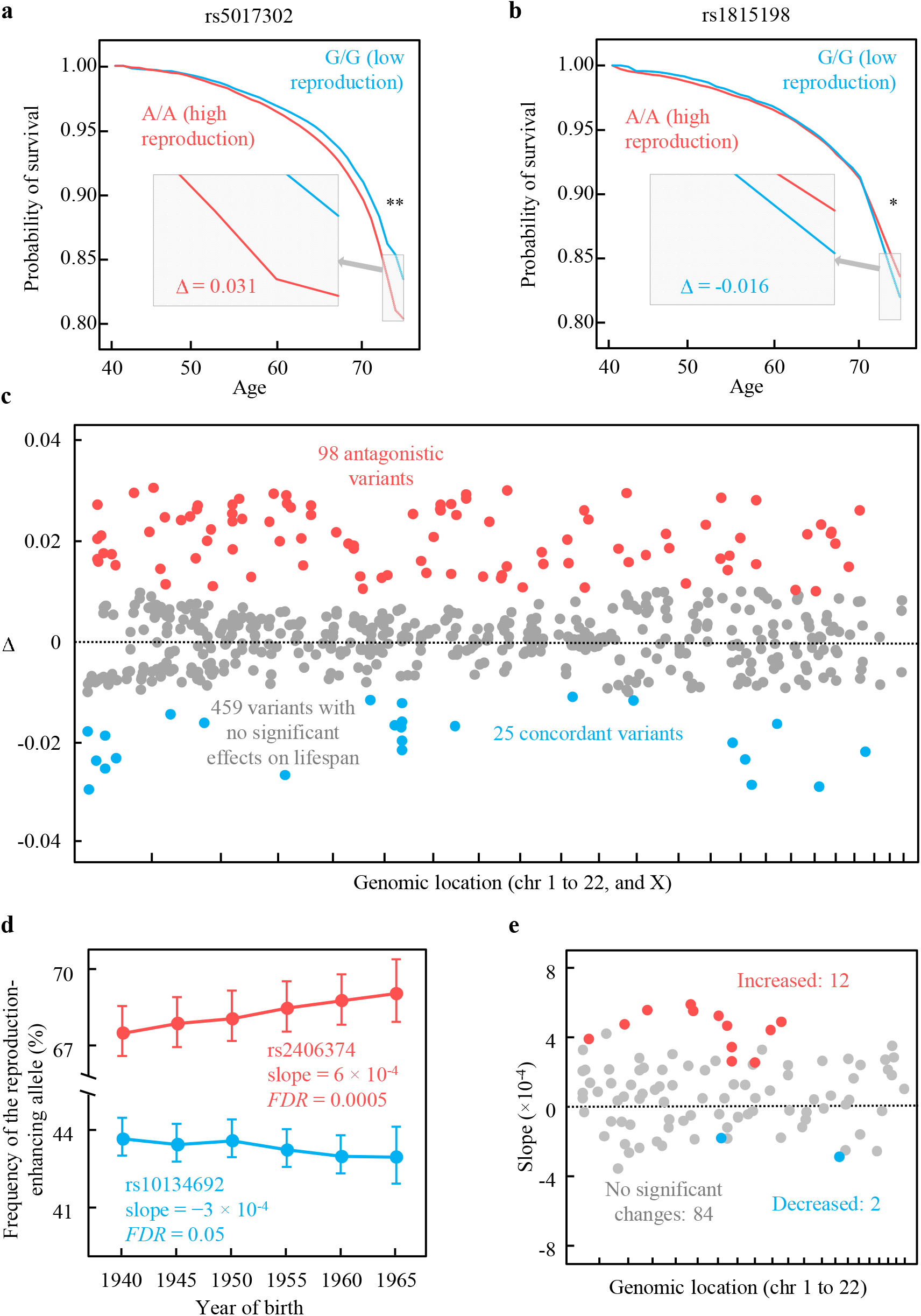
Reproduction-enhancing alleles tend to lower lifespan and be selectively favored. **a**, An example of a variant that has antagonistic effects on reproduction (nAMC) and probability of survival to age 76 (*SV*_76_). *SV*_76_ = 0.835 for G/G (lower-reproduction genotype) and 0.804 for A/A (higher-reproduction genotype) (*FDR* = 1.4×10^−6^). Δ is the difference in *SV*_76_ between lower- and higher-reproduction homozygotes for the variant considered. **b**, An example of a variant with concordant effects on reproduction (AMP) and *SV*_76_. *V*_76_ = 0.820 for T/T (lower-reproduction genotype) and 0.836 for C/C (higher-reproduction genotype) (*FDR* = 0.001). *, *FDR* < 0.05; **, *FDR* < 0.001. **c**, Effects (Δ) on *SV*_76_ of 583 reproduction-associated variants. Each dot represents one variant, whose genomic coordinate is shown on the X-axis (chromosome 1 to 22, and X for each interval from left to right). Red and blue dots indicate variants with significant antagonistic and concordant effects on *SV*_76_, respectively, whereas gray dots indicate variants with no significant effects on *SV*_76_. The horizontal line indicates no effects on *SV*_76_. **d**, Two examples of variants showing allele frequency changes over six 5-year birth cohorts (red for an increased trend and blue for a decreased trend). Slope and *FDR* are from linear regression. Error bars represent standard errors. **e**, Allele frequency changes for 98 antagonistically pleiotropic variants. Each dot represents one variant, whose genomic coordinate is shown on the X-axis (chromosome 1 to 22). Y-axis refers to the slope of the linear regression as in panel d. Red and blue dots indicate significantly positive and negative slopes, respectively, whereas gray dots indicate non-significant slopes. The horizontal line indicates a slope of 0.

Because average participants in the UK Biobank live many more years after the end of reproduction, the antagonistic pleiotropy hypothesis predicts that most mutations that increase reproduction but reduce lifespan have larger fitness advantages than disadvantages so are selectively favored. To verify this prediction, for each of the 98 variants exhibiting antagonistic pleiotropy between reproduction and lifespan, we investigated the frequency change of the allele beneficial to reproduction over the six birth cohorts after the correction for different ages of different birth cohorts at the time of the UK Biobank recruitment^25^ (see Materials and Methods). At an *FDR* of 0.05, we detected significant allele frequency increases at 12 variants and decreases at 2 variants (**Fig. 3d**), the former being significantly more prevalent than the latter (*P* = 0.006, one-tailed binomial test; **Fig. 3e, Data S4**). To account for the effect of genetic drift on allele frequency changes^26^, we applied a robustness test (see Materials and Methods) and validated 6 of the above 14 cases, all showing allele frequency increases (**Data S4**). These results demonstrate that among polymorphisms with antagonistic pleiotropy between reproduction and lifespan, alleles advantageous to reproduction tend to be selectively favored, supporting the antagonistic pleiotropy hypothesis.

### Horizontal pleiotropy between reproduction and lifespan

To understand the antagonistic pleiotropy between reproduction and lifespan, we investigated the relationship between the PGSs of the four reproductive traits and *SV*_76_ by controlling the number of children ever born (NEB = 0, 1, 2, or 3). Individuals with four or more children are too few to allow reliable estimation of *SV*_76_ in each PGS subgroup. Among individuals with the same NEB, *SV*_76_ is negatively correlated with the PGS for each of the four reproductive traits (**Fig. 4**), suggesting that the antagonistic pleiotropy between reproduction and lifespan is at least in part independent from the actual amount of reproduction. In other words, there must be horizontal antagonistic pleiotropy between reproduction and lifespan. We also performed the analysis for males and females separately and observed a negative correlation between PGS and *SV*_76_ in each stratified subgroup without significant gender-specific effects, although the negative correlation is sometimes not significant due to reduced sample sizes (**Data S5**).

**Figure 4.**
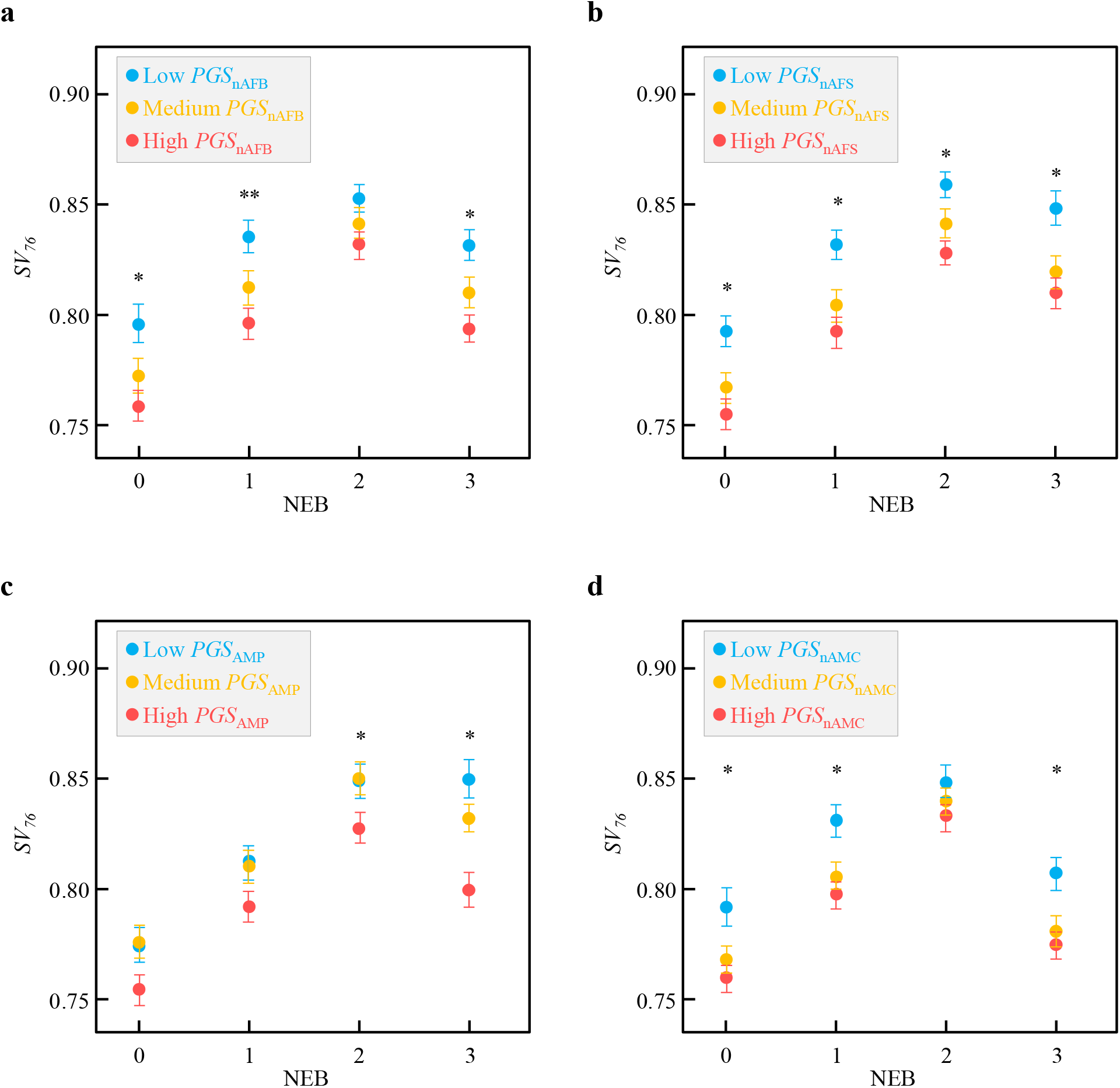
Probability of survival to age 76 (*SV*_76_) as a function of the number of children ever born (NEB) and PGSs for four reproductive traits, including negative age at first birth, nAFB (a), negative age at first sex, nAFS (b), age at menopause, AMP (c), and negative age at menarche, nAMC (d). Each dot represents individuals with the highest, medium, or lowest third of PGS considered. Error bars show 95% confidence intervals calculated by Clopper-Pearson exact methods. *, *P* < 0.05 and **, *P* < 0.001 between the low and high PGS groups. Gender-specific results are in **Data S5**.

Interestingly, given the reproductive PGS (regardless of which of the four reproductive traits is considered), individuals with NEB = 2 had the highest *SV*_76_ among the four groups of individuals with different NEBs (**Fig. 4**). Hence, when the reproductive PGS is fixed, the relationship between NEB and longevity is complex; it is positive in the low NEB range (below 2) but negative in the high NEB range (exceeding 2).

### Potential molecular mechanisms

To explore the potential mechanisms involved in the antagonistic pleiotropy between reproduction and lifespan, we queried an aggregated expression quantitative trait locus (eQTL) database—eQTL Catalogue^20^. This is because the antagonistically pleiotropic variants detected are mostly in non-coding regions and may play *cis*-regulatory roles in regulating target gene (eGene) expression. We divided the 583 reproduction-associated variants into three sets, depending on whether the variants have significant antagonistic effects on lifespan (antagonistic pleiotropy set), significant concordant effects on lifespan (concordant pleiotropy set), or no significant effects on lifespan (control set). Relative to the control set, antagonistic and concordant pleiotropy sets both contain a higher fraction of variants associated with *cis*-regulatory activities (eQTLs with empirical genome-wide significance) (**Table 2**). We defined three different types of *cis*-regulatory events: multi-context (associated with *cis*-regulatory activities in multiple tissues/cell lines), multi-eGene (associated with the expressions of multiple eGenes), and discordant (discordant effects on an eGene across tissues/cell lines). When compared with the control set, antagonistic and concordant pleiotropy sets are enriched with multi-context and multi-eGene events (**Table 2**). Additionally, the antagonistic pleiotropy set is enriched with discordant events (**Table 2**). Note that these differences are not due to potential disparities in statistical power in eQTL mapping across the three sets of variants, because we found no significant differences in minor allele frequencies across them. These results suggest that a common molecular mechanism of antagonistic pleiotropy between reproduction and lifespan is for a *cis*-regulatory mutation to influence the expressions of the same target gene in different tissues (sometimes with opposite directions) and to influence the expressions of multiple target genes.

**Table 2.**
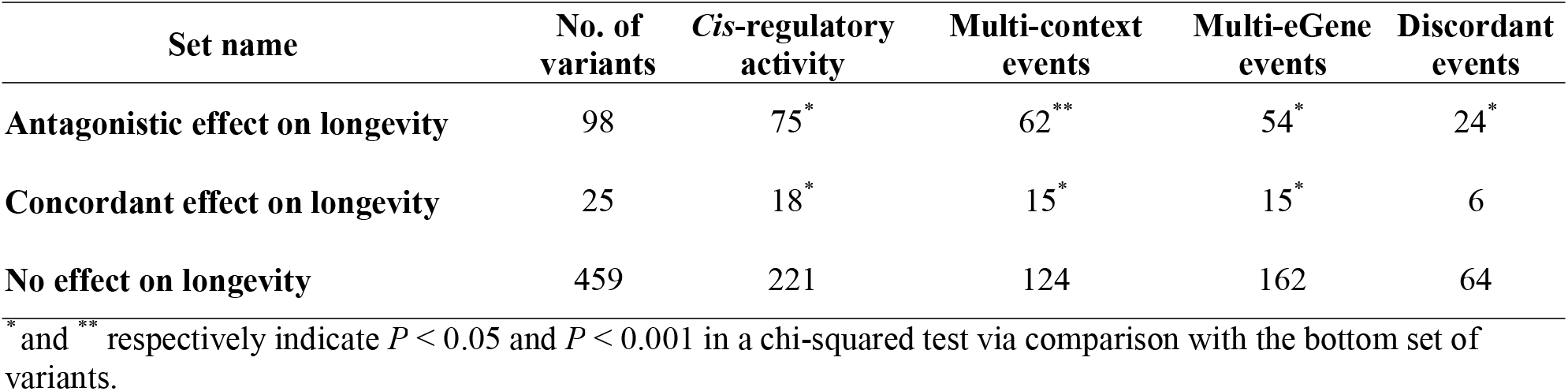
Number of reproduction-associated variants with c*is*-regulatory activities.

A total of 230 candidate eGenes from the antagonistic pleiotropy set reached empirical genome-wide significance in eQTL analysis. Significantly enriched pathways for these eGenes included those related to hormone (estrogen receptor signaling), immune response (EL-15 production), and age-related pathology (amyloid processing and protein ubiquitination pathway) (**Data S6**). Additional mechanistic insights can be gained from examining individual antagonistically pleiotropic variants. For example, variant rs12203592 at chromosome 6p25.3 has its major T allele associated with a younger age at first sexual intercourse and increased risk of mortality and shows an increase in the frequency of the T allele over 25 years (**Data S3**).

Previous GWAS suggested that the T allele is additionally associated with decreased parental lifespan^27^ and increased risk of multiple cancers such as melanoma^28^ and lung cancer^29^. This variant is also associated with multi-context, multi-eGene (*IRF4* and *EXOC2*), and discordant *cis*-regulatory events. Specifically, the T allele is correlated with higher *IRF4* expressions in the lung and whole blood but lower *IRF4* expression in skin tissues. Higher *IRF4* expressions in the human lung were reported to promote endogenous DNA damage in lung fibroblasts^29^. Another antagonistically pleiotropic variant—rs34811474 at chromosome 4p15.2—has its reproduction-enhancing G allele associated with an increased risk of osteoarthritis, an age-related degenerative disease^30^. The rs7719499-G allele, in linkage disequilibrium (*R*^2^ = 0.73, EUR population, 1000 Genome Project) with the reproduction-enhancing allele of the antagonistically pleiotropic variant rs13164856, is associated with an increased risk of cardiovascular diseases^31^. These examples show that susceptibilities to various diseases underlie the antagonistically pleiotropic effects of reproduction-enhancing alleles, consistent with findings in several previous studies^9,32,33^.

## DISCUSSION

Using the UK Biobank, we performed a series of trait-level and variant-level analyses to test the antagonistic pleiotropy hypothesis of the evolution of human aging at the genomic scale. At the trait level, we observed a strong negative genetic correlation between reproduction and parental lifespan as well as that between parental reproduction and parental lifespan, discovered that the probability of survival to the age of 76 is negatively correlated with polygenic scores for reproduction, and detected increases in these polygenic scores over 25 years. At the variant level, we found that alleles associated with higher reproduction tend to be associated with lower survival to the age 76 and that frequencies of some of these alleles have increased over the years apparently by natural selection. These findings together provide strong genome-wide evidence for the antagonistic pleiotropy hypothesis in humans. That there was ongoing selection for human reproduction as recent as a few decades ago suggests that the optimal genotype for reproduction had not been reached, possibly due to rapid environmental changes. We discovered that at least some involved antagonistically pleiotropic variants exhibit horizontal pleiotropy, meaning that they influence lifespan and reproduction through different pathways (as opposed to impacting lifespan as a result of impacting reproduction). Analysis of an eQTL database suggests that the pleiotropic effects may be often realized through *cis*-regulatory effects in multiple tissues and/or on multiple target genes.

The antagonistic pleiotropy hypothesis of aging posits that at least some mutations antagonistically affect reproduction and lifespan, without requiring such antagonistic pleiotropy be enriched. However, we found mutations impacting reproduction to be 4.9 times as likely as random, comparable mutations to influence lifespan and 7.5 times as likely to antagonistically influence lifespan. In other words, mutational antagonism between reproduction and lifespan far exceeds that expected for a random pair of traits. In terms of the potential biological reasons behind this phenomenon, it is worth mentioning the disposable soma theory of aging^34^, which posits that organisms have limited resources such that a greater investment in reproduction would lead to a lower investment in DNA repair maintenance, causing accumulation of somatic mutations and aging. The recent report of an inverse relationship between somatic mutation rate per year and lifespan across 16 mammalian species^35^ is consistent with this theory. Although this theory does not specifically invoke pleiotropy, the reproduction-lifespan tradeoff arising from resource limitation could be an explanation for antagonistic pleiotropy. Note that some mechanisms mediating the reproduction-lifespan antagonistic pleiotropy in our study are probably human-specific. For instance, a mutation that negatively impacts the educational attainment might simultaneously increase reproduction^36^ and reduce longevity^37^. By contrast, pure environmental factors that antagonistically impact reproduction and longevity (e.g., socioeconomic status) are irrelevant to the antagonistic pleiotropy hypothesis of aging, nor do they interfere with our genetic analyses such as genetic correlation and polygenic score calculations.

Given the recently raised concern on a potential publication bias toward empirical evidence for the reproduction-longevity tradeoff^38^, it is worth discussing the mutation accumulation theory of aging proposed by Peter Medawar^39^. This theory asserts that aging is an inevitable consequence of the reduction in the efficacy of natural selection with age; that is, a mutation killing youth will be strongly selected against but a lethal mutation exerting its effect only after reproduction will experience limited or even no negative selection. Over generations, late-acting deleterious mutations will accumulate, leading to an increase in mortality rates late in life. Because UK Biobank participants were at least 40 years old at the time of participation, our lifespan analysis was largely concentrated on the post-reproductive lifespan; consequently, our finding of antagonistically pleiotropic variants influencing the lifespan is consistent with the mutation accumulation theory. Nevertheless, the mutation accumulation theory posits that late-acting deleterious mutations accumulate by genetic drift while the antagonistic pleiotropy theory asserts that they are selectively favored due to the positive effects on reproduction. We found that the frequencies of reproduction-promoting alleles of some antagonistically pleiotropic variants as well as the polygenic scores for reproduction increased over years, providing strong support for the antagonistic pleiotropy theory. This, however, does not reject the mutation accumulation theory, because the two theories are not mutually exclusive; that is, the fixations of some aging-promoting alleles may follow the antagonistic pleiotropy theory (i.e., by positive selection) while others may follow the mutation accumulation theory (i.e., by genetic drift).

While our analysis focused on mutational pleiotropy between reproduction and lifespan, the relationship between reproduction and lifespan at the phenotypic level is debated. For example, a negative relationship between female fertility and longevity was reported in Chinese oldest-old individuals^40^, while an opposite trend was found in the Amish^41^. In our data, given polygenic scores for reproduction, individuals with two children have a higher probability of survival to 76 than those with 0, 1, or 3 children. Hence, the phenotypic relationship between reproduction and lifespan is complex and nonmonotonic. Furthermore, the phenotypic correlation does not equal causality because of potential confounding factors such as the socioeconomic status.

Human life expectancy, birth rate, and reproductive behavior have all changed drastically in the last few decades^42–44^. Specifically, more than half of humans live in areas of the world where birth rates have declined, along with increased incidences of contraception, abortion, and reproductive disorder^45^. The global human life expectancy at birth, on the other hand, has steadily increased from 46.5 years in 1950 to 72.8 years in 2019^46,47^. These trends of phenotypic changes are primarily driven by substantial environmental shifts including changes of lifestyles and technologies, and are opposite to the phenotypic changes caused by natural selection of the genetic variants identified in this study. This contrast indicates that, compared with environmental factors, genetic factors play a minor role in the human phenotypic changes studied here. A potential consequence of the environmental shift is that alleles with increased frequencies over time will tend to correlate with a longer lifespan, which would make our conclusion about antagonistic pleiotropy more conservative.

Our study has several limitations. First, UK Biobank participants may be a biased sample of the general population because rates of all-cause mortality and the total cancer incidence are lower in the UK Biobank than in the general population^48^. Therefore, the survival probability estimated here may be higher than that in the general population, but this factor presumably does not influence the allelic comparisons within the UK Biobank. Second, because the life expectancy of our study samples is beyond 76 years^47^, the genetic effect on lifespan is likely underestimated in our study, rendering our conclusion on the negative lifespan impact of reproduction-enhancing mutations conservative. However, if lifespan ≤ 76 years and that > 76 years are controlled by largely non-overlapping loci, our finding would be limited to the former. Third, a substantial fraction of antagonistically pleiotropic variants (23/98 = 23.5%) do not have a match in the eQTL database and we do not know how they influence reproduction and lifespan. This fraction is on par with the finding that even multiple-bulk-tissue eQTL data could not explain ~40% of the GWAS-identified variants^19,29^. Future cell-type specific eQTL and other molecular trait (e.g., methylation^49^) data might help elucidate the mechanisms by which these antagonistically pleiotropic variants act. Fourth, our study is solely based on GWAS findings and health records from European-descent individuals. Therefore, one needs to be cautious when generalizing our findings to other populations because of different social and environmental factors. Future validation of our findings in other populations, especially those currently underrepresented in biobanks^50^, is highly desired.

## MATERIALS AND METHODS

### Study participants and data used

The UK Biobank data comprise about 0.5 million participants aged 40 to 70 years, recruited between 2006 and 2010 in 22 assessment centers throughout the UK and followed up for a variety of health conditions from their recruitment date to February 17, 2016 or their date of death^17^. Participants provided a blood sample, from which DNA was extracted and genotyped using the UK BiLEVE Axiom array or Affymetrix Axiom array^17^. We used the imputed genotypes available from the UK Biobank; full details can be found in the UK Biobank imputation document (http://biobank.ctsu.ox.ac.uk/crystal/crystal/docs/impute_ukb_v1.pdf, accessed on April 16, 2022). The current study was approved by the UK Biobank (reference no. 48678), and the analyses presented were based on data from 488,377 individuals accessed through the UK Biobank (http://www.ukbiobank.ac.uk) on February 12, 2022.

From the entire set of 488,377 individuals with genotype information, we removed individuals with any of the following conditions to prevent population stratification: self-report of non-white British ethnicity, genetic principal components indicative of non-European ancestry, at least one relative in the data identified by genetic kinship, outlying level of genetic heterozygosity, and withdrawal of informed consent. Our final analysis included 276,406 unrelated individuals of European ancestry.

### Reproduction-associated variants

We collected reproduction-associated variants from NHGRI GWAS Catalog at genome-wide significance (*P* ≤ 5 × 10^−8^) by searching three types of reproductive traits: sexual maturation, reproductive behavior, and infertility risk (full lists in **Data S7**). GWASs performed or replicated in European populations were included, resulting in 891 unique variants that are associated with at least one reproductive trait.

We then tested linkage disequilibrium among these 891 variants in the EUR population (1000 Genomes Project, phase 3) by LDlink^51^. For two variants with high linkage disequilibrium (*R*^2^ > 0.8), the one with the relatively low *P* value (more significant in GWAS) was retained while the other was discarded. The variants with minor allele frequencies < 0.05 were removed. The final dataset consisted of 583 genetic variants associated with six reproductive traits: (1) nAMC, (2) AMP, (3) nAFS, (4) nAFB, (5) NEB, and (6) PCOS (**Data S1**). nAMC, nAFS, and nAFB measure the onset of reproduction, while AMP measures the end of reproduction, so together they inform the timing and potential amount of reproduction. NEB measures the actual number of children. PCOS is one of the most common causes of infertility with a relatively large genetic component^52^. By coincidence, each of these 583 variants is associated with only one of the above six reproductive traits. Hence, the direction of the allelic effect of each of these variants on reproduction was unambiguously determined.

### Estimation of survival probability

We estimated the survival probability based on Cox proportional hazard models^53^. The death records in the UK Biobank were updated quarterly with the UK National Health Service (NHS) Information Centre for participants from England and Wales and with NHS Central Register, Scotland for participants from Scotland. The latest date of death among all registered deaths in the downloaded data is 31 October 2018, and we used this date to approximate the time of last death entry and assumed that we have no mortality or viability information for the volunteers after this date. Specifically, we used five entries—age at recruitment, date of recruitment, year of birth, month of birth, and age at death—to calculate the number of individuals (*N*_i_) who are ascertained from age *i* to age *i* + 1, and the occurrence of death observed (*O*_i_) from these *N*_i_ individuals during the interval of age *i* to age *i* + 1. Using this information, we calculated the ascertained age for each individual. The death rate per year is then calculated as *h*_I_ = *O*_i_/*N*_i_ and the probability of survival to age *i* from age 40 is 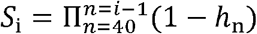. The UK Biobank data allow estimation of *h*_40_, *h*_41_, …, and *h*_75_ with *N*_i_ > 800. We estimated *h*_i_ for the homozygotes of each variant.

### Allele frequency changes over year s

We computed the frequency of the allele beneficial to reproduction at each relevant variant in each birth cohort using PLINK v2^54^. Because different birth cohorts had different ages at the time of recruitment by the UK Biobank (year 2006-2010; ~66 years old for the earliest birth cohort and ~41 years old for the latest cohort) and because some alleles affect longevity, a fair comparison of allele frequencies across birth cohorts require an equal age of participants of different cohorts. To this end, we used the probability of survival to correct the allele frequency to the age of 41 for each birth cohort. For example, let *N*_A/A_ be the number of individuals with genotype A/A in the birth cohort of 1940-1944 (age ~66 at the UK Biobank recruitment) and let *P*_66A/A_ be the probability of survival to age 66 from 41 for the genotype A/A. The number of A/A individuals at age 41 was inferred to be *N*_A/A_ divided by *P*_66A/A_, and the allele frequency was then calculated using corrected numbers of individuals of all genotypes. For each considered variant, allele frequency in a birth cohort was linearly regressed against the mid-point of the birth years of the cohort (1942.5, 1947.5, …, 1967.5), as described previously^25^. *P*-values from linear regressions were corrected for multiple testing by the Benjamini-Hochberg procedure. To consider the effect of genetic drift on allele frequency changes^26^, we employed a robustness test by further dividing individuals into 15 two-year birth cohorts (1940-1941, 1942-1943,…, 1966-1967, and 1968-1969). The allele frequency difference between the first (1940-1941) and second (1942-1943) cohort is denoted *d*_1_, and the difference between the third (1944-1945) and fourth (1946-1947) cohort is denoted *d*_2_, and so on. This resulted in seven independently estimated *d* values. We considered a variant to have changed its allele frequency consistently over the years when all seven *d* values are of the same sign, because this event has a chance probability of only 0.5^6^ = 1/64 = 0.016.

### Polygenic scores

We computed the polygenic scores for four reproductive traits (nAFB, nAFS, AMP, and nAMC) each with > 50 associated variants. All included variants were independent from one another in contributing to the trait of concern and reached genome-wide significance (see above). PLINK v2^54^ was used to calculate the polygenic score of each individual with the following formula.

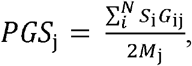

where *G*_*ij*_ is the number of reproduction-enhancing alleles at variant *i* in individual *j, S*_*i*_ is the effect size (beta) of variant *i, N* is the number of variants included in the calculation, and *M*_*i*_ is the number of non-missing variants observed in individual *j*.

### Genetic correlation surveys

We surveyed the genetic correlations computed for the significantly heritable phenotypes in the UK Biobank that are available at http://www.nealelab.is/blog/2019/10/10/genetic-correlation-results-for-heritable-phenotypes-in-the-uk-biobank. Methodological details have been published^22^.

### eQTL survey

We surveyed uniformly processed eQTLs across 69 distinct cell/tissue types from 21 available public studies aggregated by eQTL Catalogue^20^. Significant eQTLs were defined using the empirical genome-wide significance threshold as previously described^19,20^. Antagonistically pleiotropic variants were nominated with *cis*-regulatory activities if they displayed significant eQTL signals for at least one of the significant target genes (eGenes). Additionally, we defined three types of *cis*-regulatory events: (1) multi-context events: significant eQTL signals in more than one tissue/cell type, (2) multi-eGene events: significant eQTL signals for more than one eGene, and (3) discordant events: discordant directions of allelic effects for a single eGene across tissues/cell types. The nominated eGenes from antagonistically pleiotropic variants were subjected to pathway enrichment analysis by Ingenuity Pathway Analysis (IPA). Unique pathways enriched in antagonistically pleiotropic variants were defined by comparing with the enriched pathways from eGenes of reproduction-associated variants with no significant effects on lifespan (**Data S6**).

## Data availability

The following publicly available datasets were used in this work: Genetic correlations in UK Biobank, http://www.nealelab.is/blog/2019/10/10/genetic-correlation-results-for-heritable-phenotypes-in-the-uk-biobank/; GWAS Catalog, https://www.ebi.ac.uk/gwas/; eQTL Catalogue, https://www.ebi.ac.uk/eqtl/. The individual-level genotype and phenotype data were acquired from the UK Biobank (http://www.ukbiobank.ac.uk/about-biobank-uk/) under the license of Project 48678. Additional details of the data used in this work are provided in the paper and supplementary data.

## Code availability

We performed our analyses using the following publicly available software packages: LDlink, https://ldlink.nci.nih.gov/; IPA, https://www.qiagen.com/us/products/discovery-and-translational-research/next-generation-sequencing/informatics-and-data/interpretation-content-databases/ingenuity-pathway-analysis; and PLINK version 2.0: https://www.cog-genomics.org/plink/2.0/.

## ACKNOWLEDGEMENTS

We thank D. Jiang and S. Song for valuable comments. This work was supported by the research grant R35GM139484 from the U.S. National Institutes of Health to J.Z.

