## Supplementary material for "Evidence for the role of selection for reproductively advantageous alleles in human aging": Fig. S1

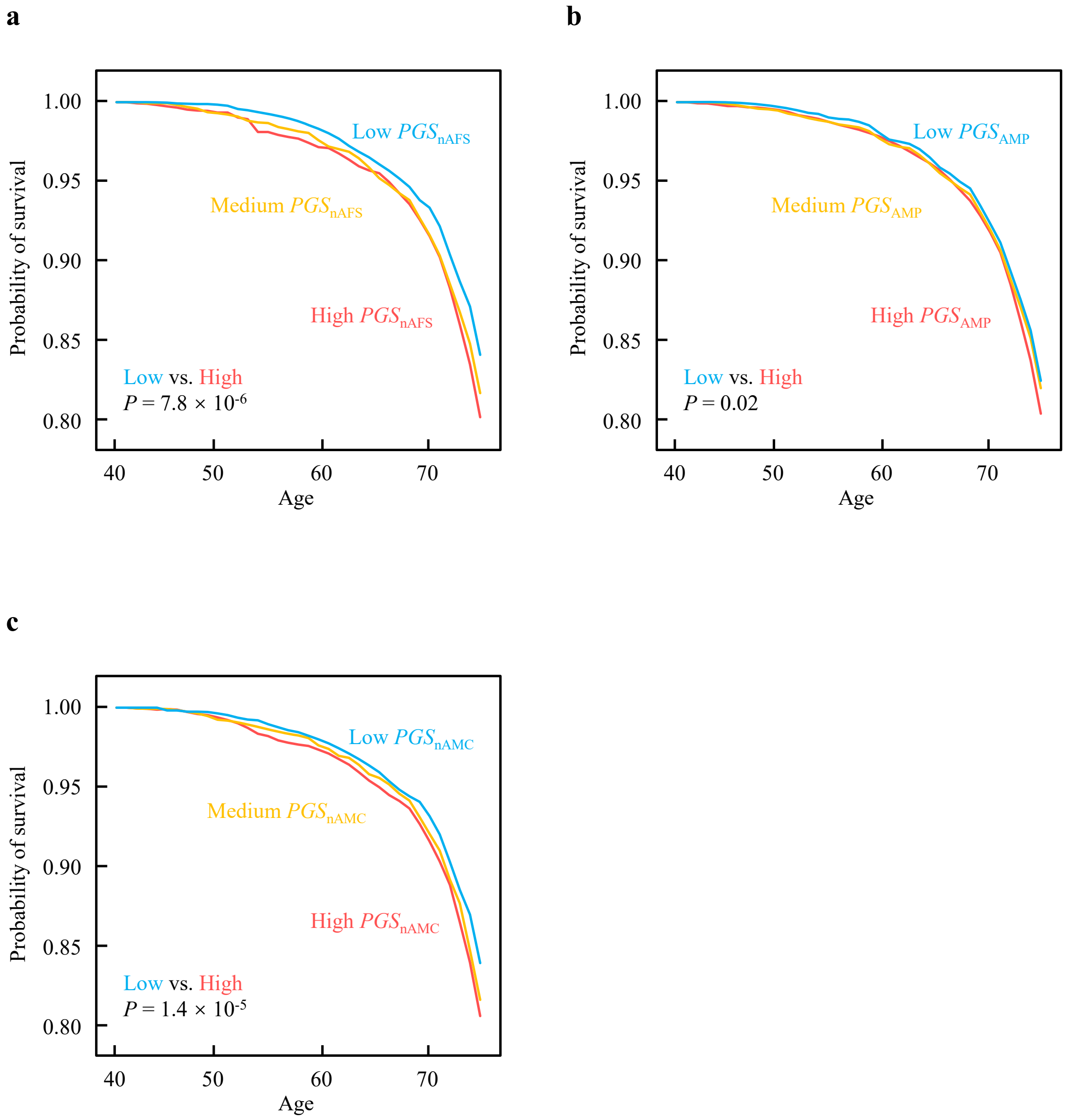


**Figure S1. Higher polygenic scores (PGSs) for the other three reproductive traits predict lower probabilities of survival to the age of 76 years ($SV_{76}$), supplementary to Figure 1a.** $SV_{76}$ of three equal-size groups of individuals with low, medium, and high PGS for negative age at first sex ($PGS_{\mathrm{nAFS}}$) (**a**), age at menopause ($PGS_{\mathrm{AMP}}$) (**b**), and negative age at menarche ($PGS_{\mathrm{nAMC}}$) (**c**).
